# Pool PaRTI: A PageRank-Based Pooling Method for Identifying Critical Residues and Enhancing Protein Sequence Representations

**DOI:** 10.1101/2024.10.04.616701

**Authors:** Alp Tartici, Gowri Nayar, Russ B Altman

## Abstract

**Motivation:** Protein language models produce token-level embeddings for each residue, resulting in an output matrix with dimensions that vary based on sequence length. However, downstream machine learning models typically require fixed-length input vectors, necessitating a pooling method to compress the output matrix into a single vector representation of the entire protein. Traditional pooling methods often result in substantial information loss, impacting downstream task performance. We aim to develop a pooling method that produces more expressive general-purpose protein embedding vectors while offering biological interpretability.

**Results:** We introduce Pool PaRTI, a novel pooling method that leverages internal transformer attention matrices and PageRank to assign token importance weights. Our unsupervised and parameter-free approach consistently prioritizes residues experimentally annotated as critical for function, assigning them higher importance scores. Across four diverse protein machine learning tasks, Pool PaRTI enables significant performance gains in predictive performance. Additionally, it enhances interpretability by identifying biologically relevant regions without relying on explicit structural data or annotated training. To assess generalizability, we evaluated Pool PaRTI with two encoder-only protein language models, confirming its robustness across different models.

**Availability and Implementation:** Pool PaRTI is implemented in Python with PyTorch and is available at https://github.com/Helix-Research-Lab/Pool_PaRTI.git. The Pool PaRTI sequence embeddings and residue importance values for all human proteins on UniProt are available at https://zenodo.org/records/15036725 for ESM2 and protBERT.

## 1 Introduction

In the rapidly evolving field of protein bioinformatics, the development of large language models for proteins (referred to as protein language models or PLMs) has marked a significant technological advancement, offering new pathways to understanding the complex nature of biological sequences [1]. Protein language models treat amino acids at different positions as distinct tokens and primarily produce embeddings for each of these tokens [1, 2]. Thus, for an input protein with (N) amino acids, the PLM generates (N) distinct token embedding vectors. These token-level representations are highly informative and useful for residue-level tasks like residue contact prediction [3] or post-translation modification prediction [4]. However, having N vectors per protein is computationally intensive and less practical for many downstream tasks at the whole sequence level, such as enzyme classification or protein-protein interaction. Therefore, many protein-level machine learning tasks require a single, fixed-length representation per protein.

To address this, traditional aggregation techniques—such as sum pooling, mean pooling, max pooling, or the direct use of the [CLS] token (a special token added to the input sequence to mark the beginning of the sequence)—offer a parameter-free way to derive a single sequence embedding. These pooling methods require no additional training or optimization, which is particularly advantageous when the training dataset for the downstream task is limited. We refer to these approaches as parameter-free task-agnostic pooling methods because they generate sequence embeddings from token embeddings without learning any task-specific parameters, resulting in embeddings that are inherently general-purpose.

Because data demands and use cases vary, we stress the distinction between parameter-free methods that produce general-purpose embeddings and parameterized methods that produce task-specific embeddings. Parameter-free methods produce general-purpose embeddings independently of any downstream task or labeled data. In contrast, parameterized methods do not modify the overall embedding generation workflow based on downstream tasks but involve some task-specific parameter optimization at the embedding generation stage. The supervision of parameter optimization comes from labeled data used for the downstream task, making the embeddings generated by these parameterized methods task-specific and not general-purpose (Figure 1).

**Figure 1:**
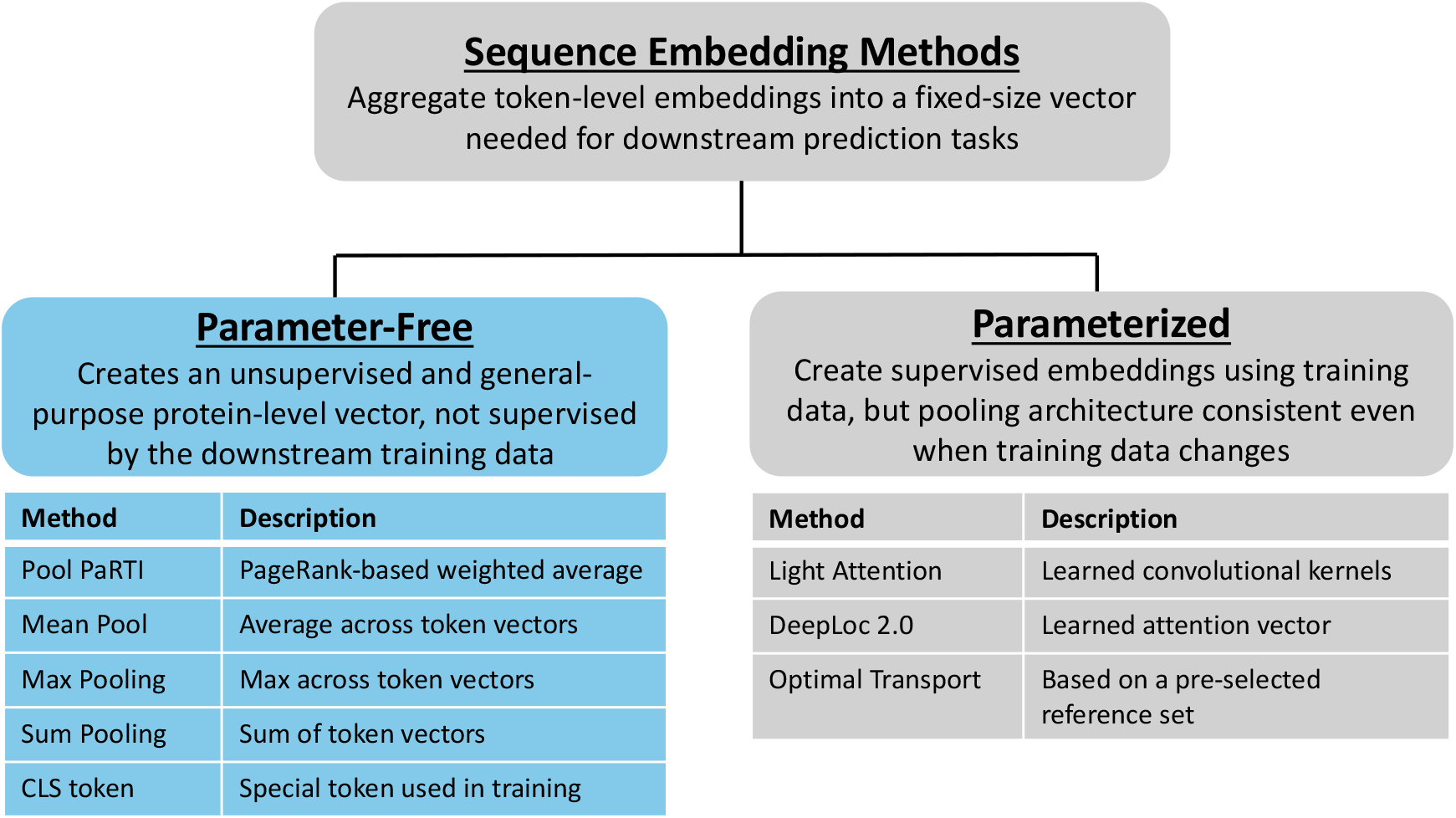
Overview of the methods used to compress token embeddings into fixed-size sequence representations for downstream protein machine learning tasks. Deep learning models require fixed-size inputs, which can be generated using either parameter-free or parameterized methods. Parameter-free methods (left) produce task-agnostic embeddings without learning task-specific parameters. Examples include Pool PaRTI, mean pooling, max pooling, CLS, and sum pooling. Parameterized methods (right) generate task-specific embeddings by learning task-dependent parameters while maintaining an architecture and workflow that can be adapted to various downstream tasks. Examples include Light Attention[9], DeepLoc2.0 [7], and optimal transport [8]

Traditional parameter-free approaches can lead to significant information loss during compression. By design, these methods may fail to capture the complex, multi-dimensional nature of biologically relevant information within the embedding, which is crucial for supporting predictive tasks in bioinformatics. Our experimental results demonstrate that protein sequence embeddings generated by these conventional parameter-free pooling methods leave room for improvement in their ability to effectively support tasks such as fold prediction, functional annotation, protein-protein interaction prediction, and subcellular localization prediction.

The potential shortfall of the traditional parameter-free pooling techniques lies in their inability to capture the variability in functional importance of segments or individual tokens. Sum pooling and mean pooling treat all tokens equally, potentially diluting the influence of critical but less frequent tokens that may be vital for specific biological functions. Conversely, max pooling may amplify noise by giving undue importance to the most extreme values, potentially skewing the representation away from biologically meaningful interpretations. In BERT-style models, due to the next sentence prediction task, the [CLS] token is instrumental in learning an aggregate summary of texts, which is pivotal for tasks such as sentiment analysis or document classification [5]. However, in protein language models like ESM2 and protBERT, the emphasis is not on predicting the continuity between two sequences but only on predicting masked amino acids through the Masked Language Model (MLM) objective function for a single sequence [2, 6]. The absence of the next sequence prediction task in these models may limit the [CLS] token’s capacity to aggregate and learn from the broader context of the sequence.

The quest for more effective pooling methods in protein language models has led to various attempts to surpass the traditional max pooling, sum pooling, mean pooling, and the [CLS] token approaches. Despite these efforts, a common limitation persists: many of these new methods are optimized for and parameterized by specific downstream tasks [7, 8, 9], which restricts the generalizability and application of the learned embeddings across different types of bioinformatics analysis. An instance is seen in the DeepLoc 2.0 model [7], which for the protein localization task, learns a scalar attention vector. This vector is used with token embeddings to assign weights in weighted average pooling, highlighting biologically significant tokens, or amino acids, that are relevant for localization [7]. However, the task-specific nature of the scalar attention vector means that it is optimized primarily for localization and may not perform well in other contexts, such as protein-protein interaction prediction. This limitation underscores a critical challenge: parameterized pooling methods offer improvements in their designated areas, but may fail adapt to the diverse needs of broader protein sequence analysis.

Adding to the diversity of pooling techniques, NaderiAlizadeh and Singh have developed an optimal transport-based pooling method that treats each token’s embedding as a sample from a probability distribution [8]. This method leverages sliced-Wasserstein distances to align these samples with a learnable reference set, generating a Euclidean embedding in the output space that encapsulates the entire protein [8]. This approach, however, relies heavily on task-specific trainable reference elements for constructing the reference probability distribution [8]. Such dependence on specific reference elements raises concerns regarding the bias that the reference selection may introduce and the ability of the generated embeddings to generalize across various tasks.

Lastly, the Light Attention method by Stärk employs a similar task-specific parameterization. Although not precisely a pooling method, Light Attention outputs a fixed-size sequence representation vector by parameterizing two sets of convolutional filters, one for learning attention values and one for the activations on the token representation matrix produced by the underlying PLM [9]. Similar to DeepLoc2.0, these convolutional filters are optimized by the labels of a downstream protein property prediction task [9], and therefore, the resulting sequence representation would depend on the task in question. The uncertainty about the adaptability of task-specific embeddings to different predictive tasks underscores a critical limitation in the existing parameterized methodologies. Moreover, the requirement to optimize parameters not only in prediction, but also in the sequence representation learning stage inevitably further increases the data demands of these methods.

The specificity of current approaches, while beneficial in narrow applications with large enough labeled data, points to a broader opportunity: the development of a parameter-free pooling method that can transcend the limitations of task-specific tuning to offer wide-reaching utility across the diverse spectrum of tasks encountered in protein bioinformatics. To our knowledge, there is no existing literature that explores the use of frozen internal parameters of the task-agnostic foundational model for developing an expressive, parameter-free pooling method to produce general-purpose protein sequence embeddings.

To address the potential limitations of the existing methods, we created a novel parameter-free pooling method that leverages the frozen internal attention matrices of transformer-based PLMs. These attention matrices, which are fundamental to the transformer architecture, contain rich information about the relationships between different tokens in the sequence [10, 1, 11]. By using this internal information, we aim to develop a more nuanced and biologically informed method for generating protein sequence embeddings.

While not directly addressing pooling methods, there is a precedence for using PLM attention maps to interpret embeddings created by these black-box models. Vig et al. [1] demonstrated significant correlations between attention patterns and well-characterized protein features, providing insights into what these models learn about protein structure and function. This approach to interpretability, although not focused on pooling, highlights the potential of leveraging internal model parameters for gaining biological insights from protein language models.

Our proposed method, Pool PaRTI (**Pool**ing by **Pa**ge**R**ank **T**oken **I**mportance), combines insights from graph theory with the internal attention mechanisms of transformer models. This approach not only aims to preserve more of the complex biological information encoded in protein sequences but also offers a task-agnostic and parameter-free solution that can be generalized to various protein language models and downstream tasks.

In this paper, we present Pool PaRTI and demonstrate its effectiveness in sequence representation across four diverse protein-related tasks. We show that with two different underlying PLMs, the protein residues that the Pool PaRTI workflow highlights align with residues that have been experimentally annotated as critical for activity. We explore biologically interpretable patterns identified by the Pool PaRTI workflow, visually demonstrating its ability to detect functionally relevant regions in proteins without explicit structural or functional training. Furthermore, we provide empirical evidence for the performance gains with Pool PaRTI compared to traditional parameter-free sequence embedding generation methods, namely sum pooling, mean pooling, max pooling, and the [CLS] token . This work represents a significant step towards more effective and biologically meaningful protein sequence representations, with potential implications for enhancing a wide range of bioinformatics discovery applications from protein functional annotation to variant effect prediction and possibly beyond to other domains of sequence analysis in machine learning.

## 2 Systems and Methods

Pool PaRTI capitalizes on the internal attention matrices of transformer-based protein language models. These matrices provide a rich representation of the relationships between tokens in the sequence, offering a more nuanced view of token interactions than traditional pooling methods.

Pool PaRTI uses a weighted average of token embeddings to construct a comprehensive sequence embedding vector. The weight assignment process involves leveraging the attention matrices across all layers of the transformer model. Unlike raw attention heads that only yield pairwise token importance values, Pool PaRTI conceptualizes the protein attention matrices as adjacency matrices in a directed and weighted graph model. To generate the source matrix for our graph, we perform pixel-wise max pooling across all layers of the internal attention matrices. This operation captures the strongest attention relationships across the entire model. Furthermore, previous work has shown that attention heads, not only in the final layer but also in intermediate layers, can highlight regions with functionally important protein domains [11]. Within the resulting graph, amino acid tokens are represented as nodes, and their interactions, determined by the pooled attention from the query to the key, serve as directed edges with corresponding weights (Figure 2). Upon constructing this graph, we run the PageRank algorithm on the graph to extract normalized node importance values, which correspond directly to token weights in weighted average pooling (Algorithm 1). We normalize the token weights to sum to one to ensure that the output sequence embedding is independent of the input sequence length.

**Figure 2:**
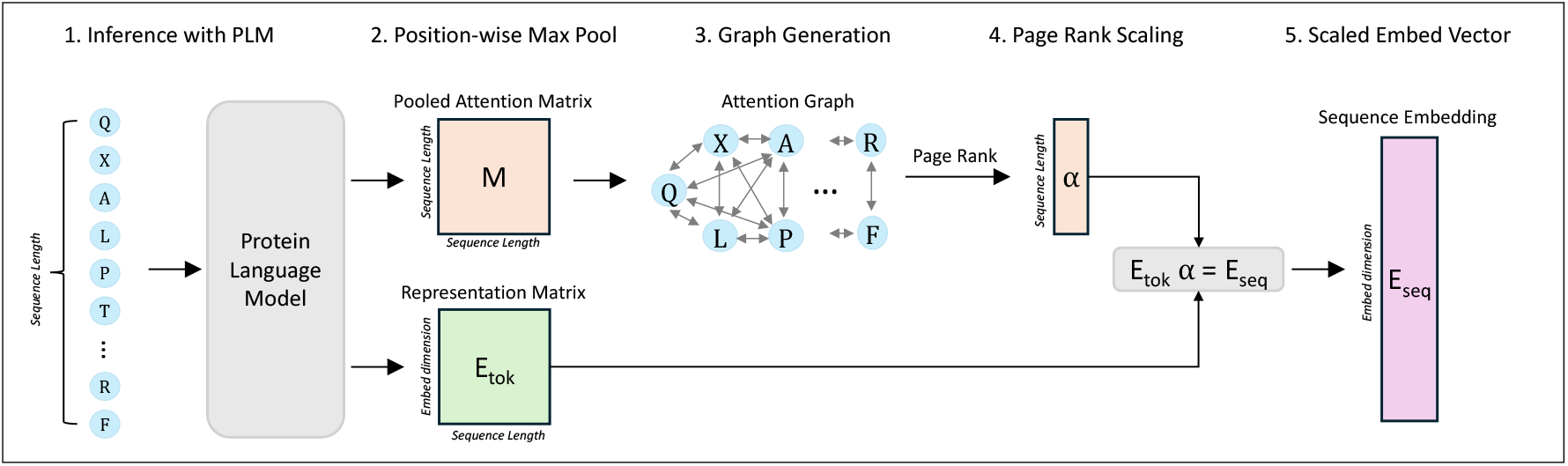
Graphical Summary of the Pool PaRTI Algorithm. **Step (1)** is feeding the whole sequence into an encoder PLM and extracting representation and attention matrices. **Step (2)** is position-wise max pooling of all attention matrices into a single matrix. **Step (3)** is treating the pooled attention matrix like an adjacency matrix and converting it into a fully connected directed and weighted graph G, where nodes are tokens and edges are attention values. **Step (4)** is applying PageRank on graph G to extract token (or node) importance weight values *α*. **Step (5)** is computing the output sequence embedding by taking the importance weighted average of residue embeddings.

Pool PaRTI algorithm is run only once per protein in inference time and not run during the training phase. We selected the PageRank algorithm for its effectiveness in graphs with weighted and directed edges, useful for delineating token significance in the complex hierarchy within protein sequences [12]. For a graph *G* with *N* nodes and *E* edges, the PageRank algorithm with a random surfer model has a space complexity of *O*(*N* + *E*) [12] because at each iteration, the algorithm only stores the edge connectivity, the weights, if any, and the current node importance values. This level of space complexity is manageable within the context of the much larger space used by the foundation model itself. At each iteration, PageRank computes importance transfer along all the edges. Hence, the time complexity is *O*(*kE*) for convergence over k iterations [12]. Given that the attention matrix from BERT style models typically results in a fully connected graph where *E* grows quadratically with *N*, the computational cost of applying PageRank directly on the original matrix would scale quadratically with *N*, the length of the protein sequence.

## 3 Experiments and Results

We conducted a series of experiments to evaluate the performance of Pool PaRTI across four different protein-related tasks. For demonstrations, we used ESM2 650M [2] and protBERT [6] protein language models to generate token embeddings and extract attention matrices due to their manageable size and all-to-all attention-based encoder architectures.

### 3.1 Importance weights attributed to residues are reproducible and align with experimental annotations

We investigated how Pool PaRTI assigns importance weights to protein residues and whether the designation of the highest-importance residues depends on the choice of PLM. To assess the level of general alignment, we computed Pool PaRTI importance scores separately using attention matrices from both ESM2 and protBERT. We then quantified the Jaccard similarity between the top *k* percentile of important residues for various *k* values. Our results show that the agreement between ESM2-based and protBERT-based importance weights is significantly higher than expected under both a theoretical random weight assignment model (detailed in Appendix A.1) and an empirical null background (Figure 3). The latter compares the top *k* percent of important residues from randomly chosen proteins of the same length. These comparisons suggest that Pool PaRTI’s importance weight assignment is reproducible and not due to random chance or regional weight assignment bias.

**Figure 3:**
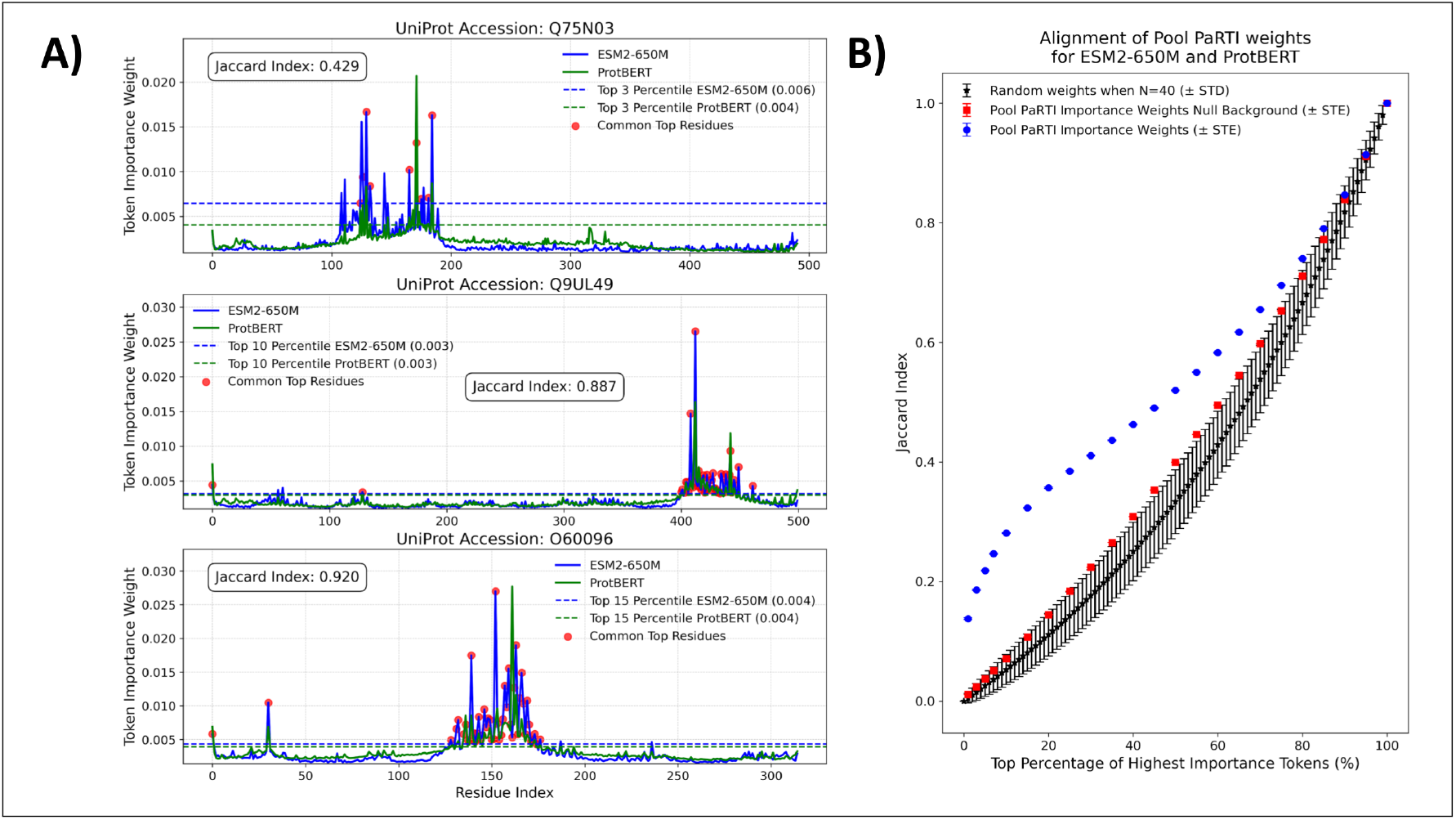
A) The examples of alignments between the Pool PaRTI token importance weights assigned based on ESM2 and protBERT inputs, with the Jaccard indices calculated for illustration at three different percentile cutoffs (3, 10, and 20). B) The statistical summary of the aligment between Pool PaRTI embeddings from two different PLMS, compared to the theoretical and empirical distributions of random weight assignments.

#### Algorithm 1

Pool PaRTI: Protein Sequence Embedding Calculation

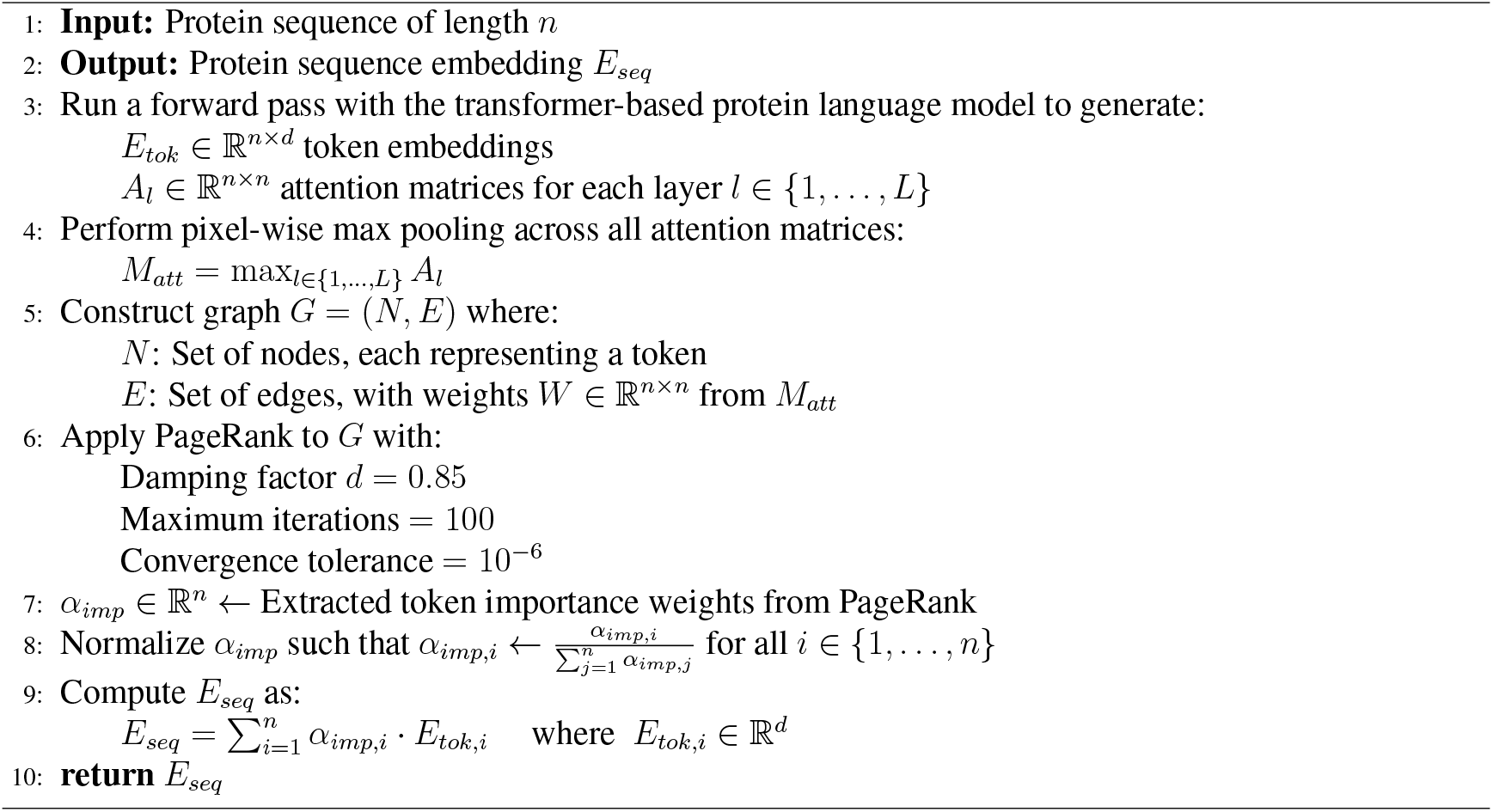

We further examined whether residues assigned high importance by Pool PaRTI overlap with experimentally validated functional residues from the Catalytic Site Atlas [13]. We found that functionally annotated residues are disproportionately ranked among the most important positions in both ESM2- and protBERT-derived importance scores (Figure 4). This demonstrates that Pool PaRTI not only generalizes across different PLMs but also effectively highlights critical residues without explicit structural supervision.

**Figure 4:**
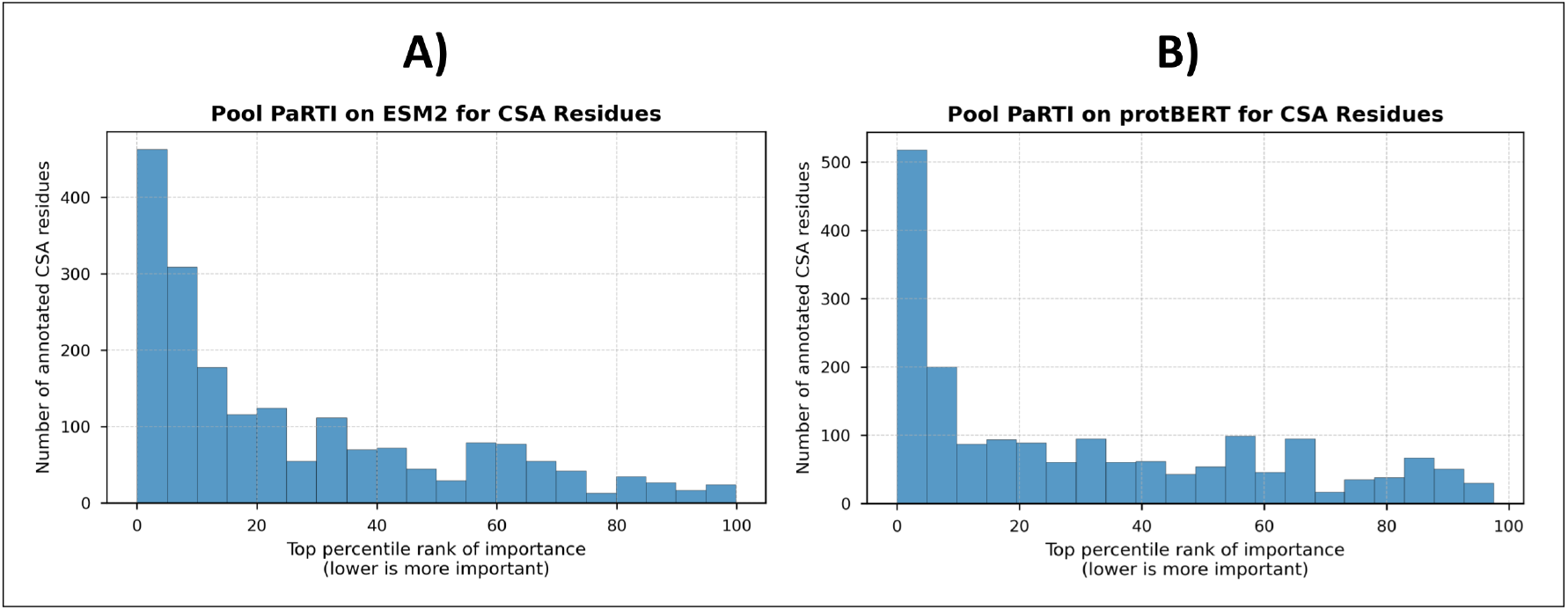
Percentile rank distribution of residues experimentally annotated as functionally important in the Catalytic Site Atlas, based on Pool PaRTI importance score assignments for A) ESM2 and B) protBERT inputs. The clear non-uniform distribution and heavy skew towards toward the top percentiles demonstrate Pool PaRTI’s ability to assign higher importance weights to critical residues.

To investigate the biological relevance of the importance weights assigned by our method, we conducted an in-depth analysis of several well-characterized proteins. This examination revealed a striking visual overlap between residues assigned higher importance weights by Pool PaRTI and known functionally or structurally significant regions within these proteins. A notable example is observed in Cytochrome C, a crucial component of the electron transport chain [14]. Pool PaRTI assigned higher importance weights to residues within the heme-binding regions, which are essential for the protein’s electron transfer function [14] (Figure 5A). Similarly, in superoxide dismutase-1 (SOD1), an enzyme critical for cellular antioxidant defense [15], our method highlighted regions proximal to the zinc-binding domain (Figure 5B). This domain is integral to SOD1’s catalytic activity and structural stability, and mutations therein are implicated in Amyothropic Lateral Sclerosis (ALS) [16]. Taken together, these findings are particularly significant as they demonstrate Pool PaRTI’s capacity to identify biologically relevant regions without explicit training on structural or functional data. This emergent property suggests that our method captures intrinsic sequence features that correlate with functional importance, potentially offering new insights into protein structure-function relationships.

**Figure 5:**
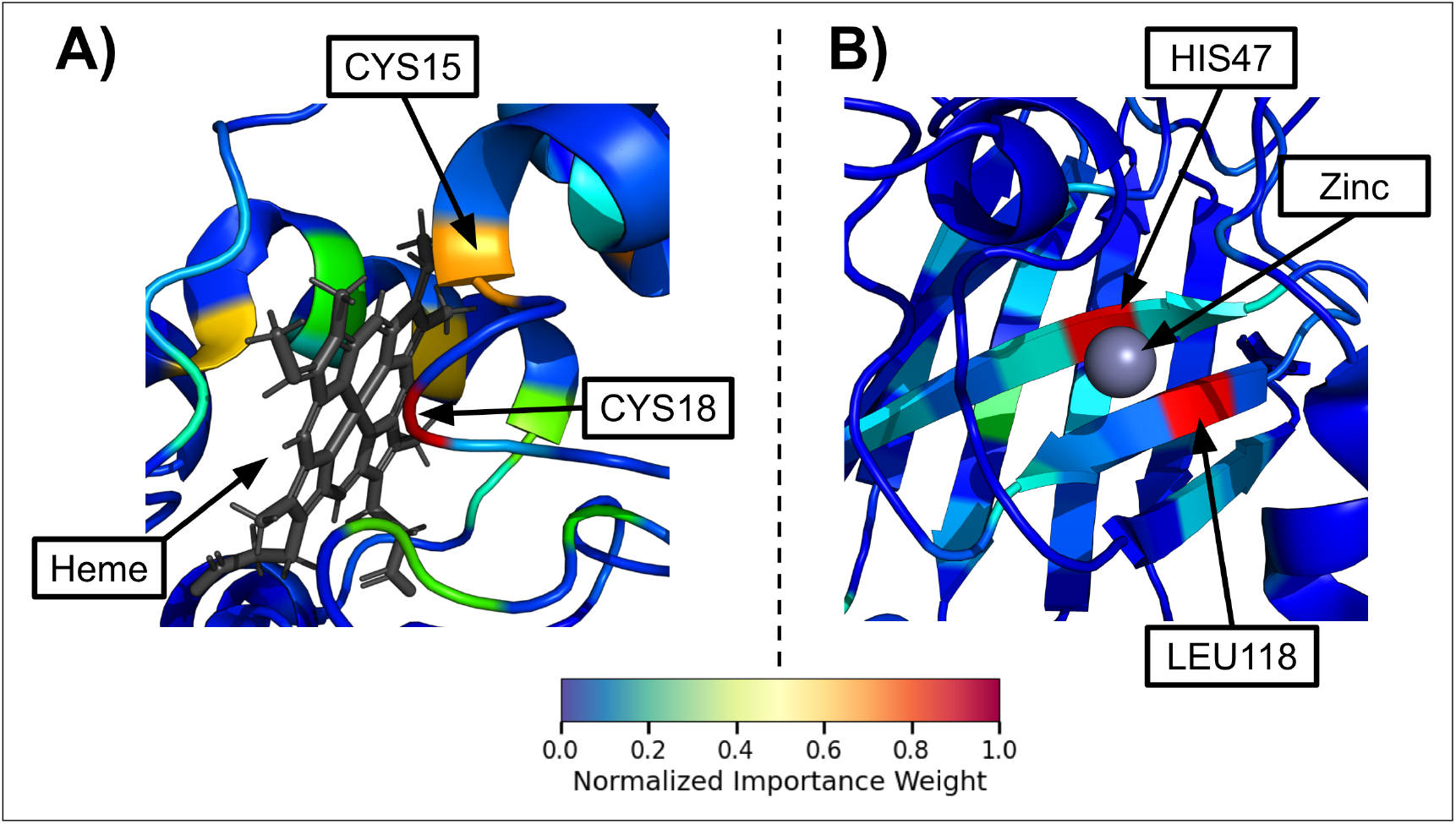
Visual demonstrations of the unsupervised importance weights assigned by Pool PaRTI to protein residues. **A)** In Cytochrome C (105 residues), Pool PaRTI ranks CYS18 and CYS15 as the first and third most important residues, respectively. These residues are essential for the protein’s Heme-binding activity. **B)** In Superoxide Dismutase-1 (154 residues), Pool PaRTI identifies HIS47 and LEU118 as the most and second-most important residues, respectively, both of which play a crucial role in Zinc binding.

**Figure 6:**
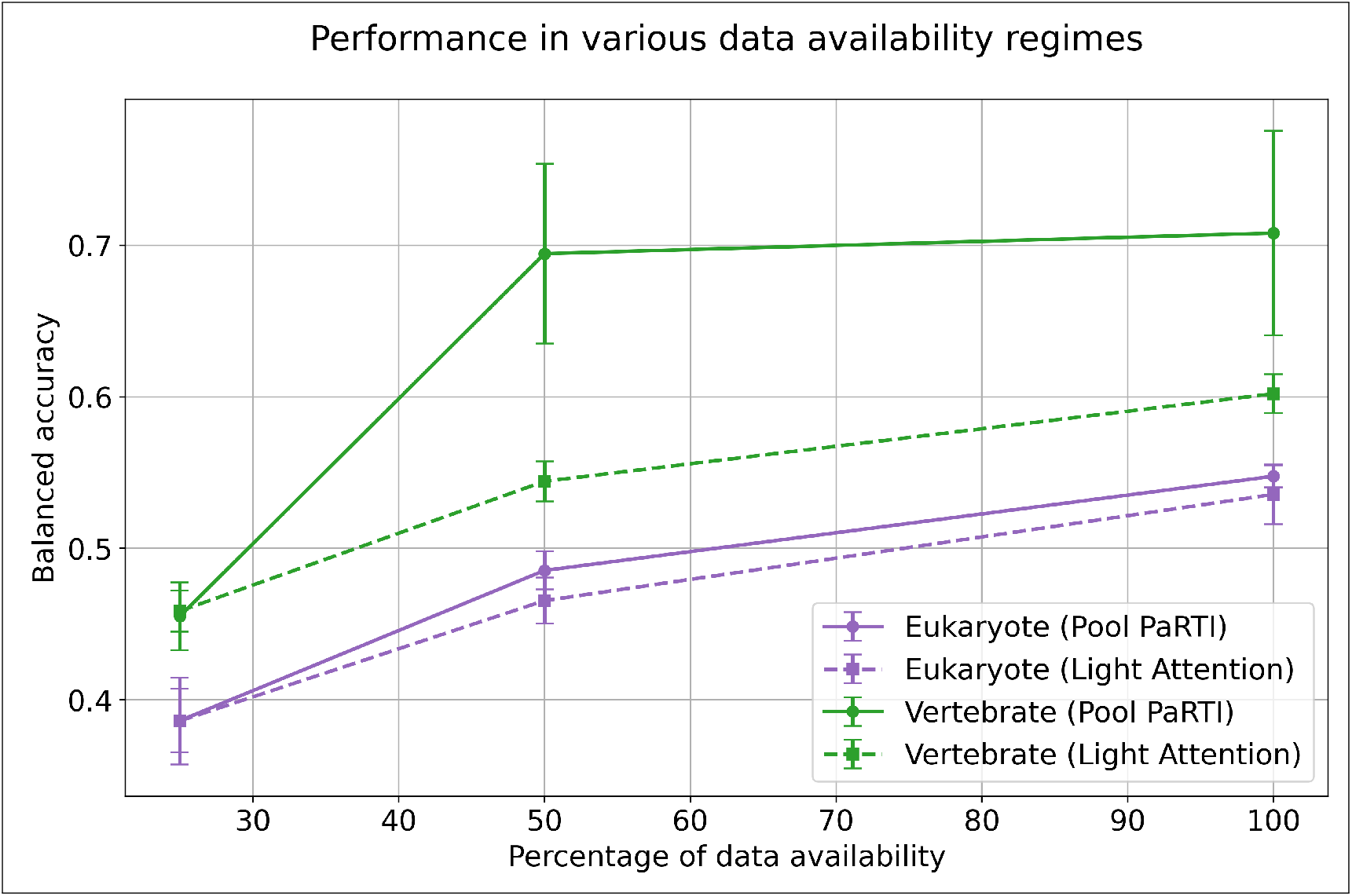
The performance of Pool PaRTI-based predictor versus the parameter-heavy Light Attention model in predicting the Level 1 EC classes in various degrees of training data scarcity.

### 3.2 Pool PaRTI embeddings improve performance in downstream prediction tasks

Having found that Pool PaRTI can discern the hierarchy of critical residues from the pairwise attention values, we tested whether Pool PaRTI embeddings generated based on this hierarchy can improve predictive performances in downstream protein sequence level bioinformatics tasks. We validated Pool PaRTI’s sequence embedding quality across four distinct protein-related prediction tasks— SCOP protein fold classification and enzyme class prediction through direct embedding retrieval, followed by protein-protein interaction predictions and multilabel subcellular localization using simple feed-forward neural networks. We used consistent model architectures, computational resources, and embedding retrieval schemes across the competing pooling methods. We designed each task to assess Pool PaRTI’s efficacy in retaining the information content of raw token embedding matrices compared to the baselines of four traditional parameter-free methods: [CLS] token, mean pooling, sum pooling, and max pooling.

The methodological consistency in the downstream tasks (e.g., model sizes, hyperparameter tuning scheme) ensures that the observed performance enhancements are directly attributable to our pooling method. To ensure the statistical significance of our results, we repeatedly sampled balanced subsets of the dataset and averaged across the samples. We conducted the neural network-based experiments using 10 random seeds for all methods under comparison after finding the ideal hyperparameter sets for each competing pooling method. See Appendix A.2 for the description of the relevant metrics for each task.

In the first experiment, we evaluated how well the sequence embeddings generated by different pooling methods align with SCOP (Structural Classification of Proteins) families. SCOP is a database that classifies protein structures hierarchically, with level 2 corresponding to the fold class [17, 18]. For each query protein, we retrieved neighboring proteins with the highest cosine similarity in sequence embedding. For both ESM2 and protBERT as the underlying PLM, Pool PaRTI achieved higher scores in both retrieval metrics compared to all baseline methods, indicating better preservation of information related to protein structure (Table 1 and 2).

**Table 1:** The performances in Embedding-based Retrieval of SCOP Level 2 Fold Categories averaged over 50 class-balanced sampling rounds using ESM2.

| Metric | Pooling |  |  |  |  |
| --- | --- | --- | --- | --- | --- |
|  | [CLS] token | Mean pooling | Max pooling | Sum pooling | Pool PaRTI |
| Precision@10% | 0.496 | 0.474 | 0.500 | 0.476 | **0.508** |
| Mean Reciprocal Rank | 0.667 | 0.657 | 0.674 | 0.658 | **0.682** |

The standard error values are all below 0.0005 for the reported metrics as a consequence of repeated subsampling.
\*\*\*p < 0.01, \*\*p < 0.05, \*p < 0.10. p values correspond to the null hypothesis that the respective pooling method has the same performance as Pool PaRTI, tested by Mann-Whitney U test.

**Table 2:** The performances in Embedding-based Retrieval of SCOP Level 2 Fold Categories averaged over 50 class-balanced sampling rounds using protBERT.

| Metric | Pooling |  |  |  |  |
| --- | --- | --- | --- | --- | --- |
|  | [CLS] token | Mean pooling | Max pooling | Sum pooling | Pool PaRTI |
| Precision@10% | 0.248 | 0.326 | 0.294 | 0.325 | **0.342** |
| Mean Reciprocal Rank | 0.461 | 0.533 | 0.504 | 0.532 | **0.549** |

The standard error values are all below 0.0005 for the reported metrics as a consequence of repeated subsampling.
\*\*\*p < 0.01, \*\*p < 0.05, \*p < 0.10. p values correspond to the null hypothesis that the respective pooling method has the same performance as Pool PaRTI, tested by Mann-Whitney U test.

The second experiment evaluates the alignment of top-level Enzyme Commission (EC) number of proteins with the embeddings generated by different methods. We used the gold standard ECPred40 dataset from [19]. EC numbers were chosen over Gene Ontology (GO) codes due to their robustness; GO term prediction suffers from significant redundancy and label noise [20]. When retrieving the enzyme class, Pool PaRTI outperforms the baseline methods in both retrieval metrics when ESM2 is used as the underlying PLM (Table 3). However, with protBERT as the underlying PLM, Pool PaRTI ranks first in mean reciprocal rank but third in precision at 10% of the class size, closely behind mean pooling and sum pooling (Table 4).

**Table 3:** The performances in Embedding-based Retrieval of EC Level 1 enzyme category averaged over multiple class-balanced sampling rounds using ESM2.

| Metric | Pooling |  |  |  |  |
| --- | --- | --- | --- | --- | --- |
|  | [CLS] token | Mean pooling | Max pooling | Sum pooling | Pool PaRTI |
| Precision@10% | 0.373 | 0.420 | 0.421 | 0.422 | **0.429** |
| Mean Reciprocal Rank | 0.663 | 0.701 | 0.698 | 0.698 | **0.710** |

The standard error values are all below 0.0005 for the reported metrics as a consequence of repeated subsampling.
\*\*\*p < 0.01, \*\*p < 0.05, \*p < 0.10. p values correspond to the null hypothesis that the respective pooling method has the same performance as Pool PaRTI, tested by Mann-Whitney U test.

**Table 4:** The performances in Embedding-based Retrieval of EC Level 1 enzyme category averaged over multiple class-balanced sampling rounds using protBERT.

| Metric | Pooling |  |  |  |  |
| --- | --- | --- | --- | --- | --- |
|  | [CLS] token | Mean pooling | Max pooling | Sum pooling | Pool PaRTI |
| Precision@10% | 0.238 | **0.283** | 0.268 | **0.283** | 0.276 |
| Mean Reciprocal Rank | 0.569 | 0.616 | 0.604 | 0.617 | **0.625** |

The standard error values are all below 0.0005 for the reported metrics as a consequence of repeated subsampling.
\*\*\*p < 0.01, \*\*p < 0.05, \*p < 0.10. p values correspond to the null hypothesis that the respective pooling method has the same performance as Pool PaRTI, tested by Mann-Whitney U test.

Our third task tests performance on predicting whether two given protein sequences interact one another, using gold standard data split taken from [21]. This binary classification task evaluates the methods’ ability to retain the expressive power regarding the complex biological interactions across sequences. Unlike the earlier experiments, this task takes two input embeddings to predict a single label. Therefore we opted for developing a small-scale neural network model instead of using direct embedding retrieval for prediction. When ESM2 is the underlying PLM, Pool PaRTI significantly outperforms the [CLS] token, sum pooling, and max pooling methods in terms of accuracy, MCC, and AUPRC. Pool PaRTI outperforms mean pooling as well on all three metrics, albeit with less statistical significance (Table 5). When protBERT is the underlying PLM, Pool PaRTI ties for the first place in AUPRC while closely following mean pooling and sum pooling in accuracy and MCC 6.

**Table 5:** The performances in protein-protein interaction prediction averaged over 10 random seeds using ESM2.

| Metric | Pooling |  |  |  |  |
| --- | --- | --- | --- | --- | --- |
|  | [CLS] token | Mean pooling | Max pooling | Sum pooling | Pool PaRTI |
| Accuracy | 0.627±0.004*** | 0.642±0.002* | 0.591±0.002*** | 0.627±0.002*** | **0.646±0.002** |
| MCC | 0.260±0.006*** | 0.285±0.003* | 0.186±0.003*** | 0.254±0.003*** | **0.291±0.005** |
| AUPRC | 0.686±0.003*** | 0.697±0.002 | 0.628±0.002*** | 0.671±0.003*** | **0.699±0.002** |

\*\*\*p < 0.01, \*\*p < 0.05, \*p < 0.10. p values correspond to the null hypothesis that the respective pooling method has the same performance as Pool PaRTI, tested by Mann-Whitney U test.

**Table 6:** The performances in protein-protein interaction prediction averaged over 10 random seeds using protBERT.

| Metric | Pooling |  |  |  |  |
| --- | --- | --- | --- | --- | --- |
|  | [CLS] token | Mean pooling | Max pooling | Sum pooling | Pool PaRTI |
| Accuracy | 0.583±0.001*** | 0.641±0.001 | 0.598±0.001*** | **0.642±0.001\*** | 0.639±0.001 |
| MCC | 0.167±0.002*** | **0.286±0.001\*\*** | 0.197±0.001*** | 0.282±0.001 | 0.279±0.002 |
| AUPRC | 0.625±0.001*** | **0.701±0.002** | 0.642±0.001*** | 0.700±0.001 | **0.701±0.001** |

\*\*\*p < 0.01, \*\*p < 0.05, \*p < 0.10. p values correspond to the null hypothesis that the respective pooling method has the same performance as Pool PaRTI, tested by Mann-Whitney U test.

In our fourth downstream experiment, we aimed to determine if a given protein sequence can be classified into any of eight possible subcellular locations, such as the nucleus, cell membrane, mitochondria, or endoplasmic reticulum, using the dataset split in [7]. This task is inherently a multilabel binary classification since a protein can localize to multiple cellular compartments simultaneously. Therefore, we chose to develop neural network classifiers to predict the labels. For both ESM2 and protBERT, Pool PaRTI significantly outperforms all four baseline methods in terms of accuracy, Matthews Correlation Coefficient (MCC) and Area Under the Precision-Recall Curve (AUPRC) metrics. Furthermore, it significantly outperforms all four baselines in Jaccard index, indicating its robust capability in extracting spatial information and handling the complexities of multilabel classification (Tables 7, 8).

**Table 7:** The performances in subcellular localization prediction task averaged over 10 random seeds using ESM2.

| Metric | Pooling |  |  |  |  |
| --- | --- | --- | --- | --- | --- |
|  | [CLS] token | Mean pooling | Max pooling | Sum pooling | Pool PaRTI |
| Acc. avg | 0.872±0.004*** | 0.873±0.000*** | 0.843±0.004*** | 0.772±0.027*** | **0.905±0.001** |
| MCC avg | 0.256±0.033*** | 0.306±0.004*** | 0.198±0.037*** | 0.082±0.052*** | **0.572±0.005** |
| AUPRC avg | 0.330±0.030*** | 0.415±0.009*** | 0.195±0.023*** | 0.158±0.003*** | **0.493±0.003** |
| Jaccard index | 0.450±0.059*** | 0.520±0.003*** | 0.239±0.030*** | 0.245±0.043*** | **0.639±0.002** |

\*\*\*p < 0.01, \*\*p < 0.05, \*p < 0.10. p values correspond to the null hypothesis that the respective pooling method has the same performance as Pool PaRTI, tested by Mann-Whitney U test.

**Table 8:** The performances in subcellular localization prediction task averaged over 10 random seeds using protBERT.

| Metric | Pooling |  |  |  |  |
| --- | --- | --- | --- | --- | --- |
|  | [CLS] token | Mean pooling | Max pooling | Sum pooling | Pool PaRTI |
| Acc. avg | 0.793±0.013 | 0.792±0.006 | 0.738±0.008*** | 0.766±0.006*** | **0.798±0.006** |
| MCC avg. | 0.360±0.006*** | 0.406±0.005 | 0.317±0.003*** | 0.337±0.004*** | **0.410±0.005** |
| AUPRC avg | 0.423±0.004*** | 0.487±0.004 | 0.427±0.003*** | 0.380±0.003*** | **0.495±0.003** |
| Jaccard index | 0.405±0.012 | 0.407±0.008 | 0.351±0.004*** | 0.364±0.006*** | **0.414±0.006** |

\*\*\*p < 0.01, \*\*p < 0.05, \*p < 0.10. p values correspond to the null hypothesis that the respective pooling method has the same performance as Pool PaRTI, tested by Mann-Whitney U test.

As shown in Tables 1-8, Pool PaRTI’s closest competitors in head-to-head comparisons are mean pooling and sum pooling. When ESM2 is used as the underlying PLM, Pool PaRTI achieves significant performance gains across all four tasks, outperforming all other embedding generation methods. However, with protBERT as the underlying PLM, its advantage diminishes. In this case, Pool PaRTI leads in two of the four tasks while remaining competitive with mean pooling and sum pooling in the other two. Regardless of the underlying PLM, Pool PaRTI consistently outperforms max pooling and the [CLS] token across all tasks and metrics.

Beyond comparisons with parameter-free sequence embedding methods, we evaluated Pool PaRTI against a heavily parameterized approach, Light Attention, in a scenario with limited labeled data. Using the ECPred40 dataset [19], we predicted the top-level EC class, training models on human proteins and testing them separately on unseen vertebrate and eukaryotic proteins. Vertebrate proteins, a subset of eukaryote proteins, are more similar to the training data. To ensure a fair comparison, we kept the prediction head model size fixed at 1.5 million parameters. However, Light Attention required an additional 29.5 million parameters to optimize convolutional kernels for generating fixed-size (1280-dimensional) sequence embeddings, whereas Pool PaRTI remained entirely parameter-free in embedding generation.

We hypothesized that Light Attention’s reliance on parameterized convolutional kernels would make it significantly more data-hungry, giving Pool PaRTI an advantage when training data is limited. The original training and validation sets contained 1,752 human proteins. To evaluate performance degradation as data availability decreased, we optimized Pool PaRTI-based and Light Attention-based models separately at 100%, 50%, and 25% of the dataset, ensuring that smaller datasets still maintained class balance. In predicting enzymatic activity for vertebrate proteins, the Pool PaRTI-based model outperformed the Light Attention model at 50% and 100% data availability and converged to the same accuracy as Light Attention at 25%.

To assess computational efficiency, we measured the runtime of Pool PaRTI on a CPU (Intel(R) Xeon(R) Gold 5118 @ 2.30GHz) as a function of sequence length, confirming its quadratic time complexity in sequence length (Figure 7, Appendix A.3). Our empirical analysis demonstrates that Pool PaRTI does not introduce prohibitive computational overhead compared to other pooling methods at input sizes relevant to the human proteome. Detailed model configurations for all four experiments are provided in Appendix A.4.

**Figure 7:**
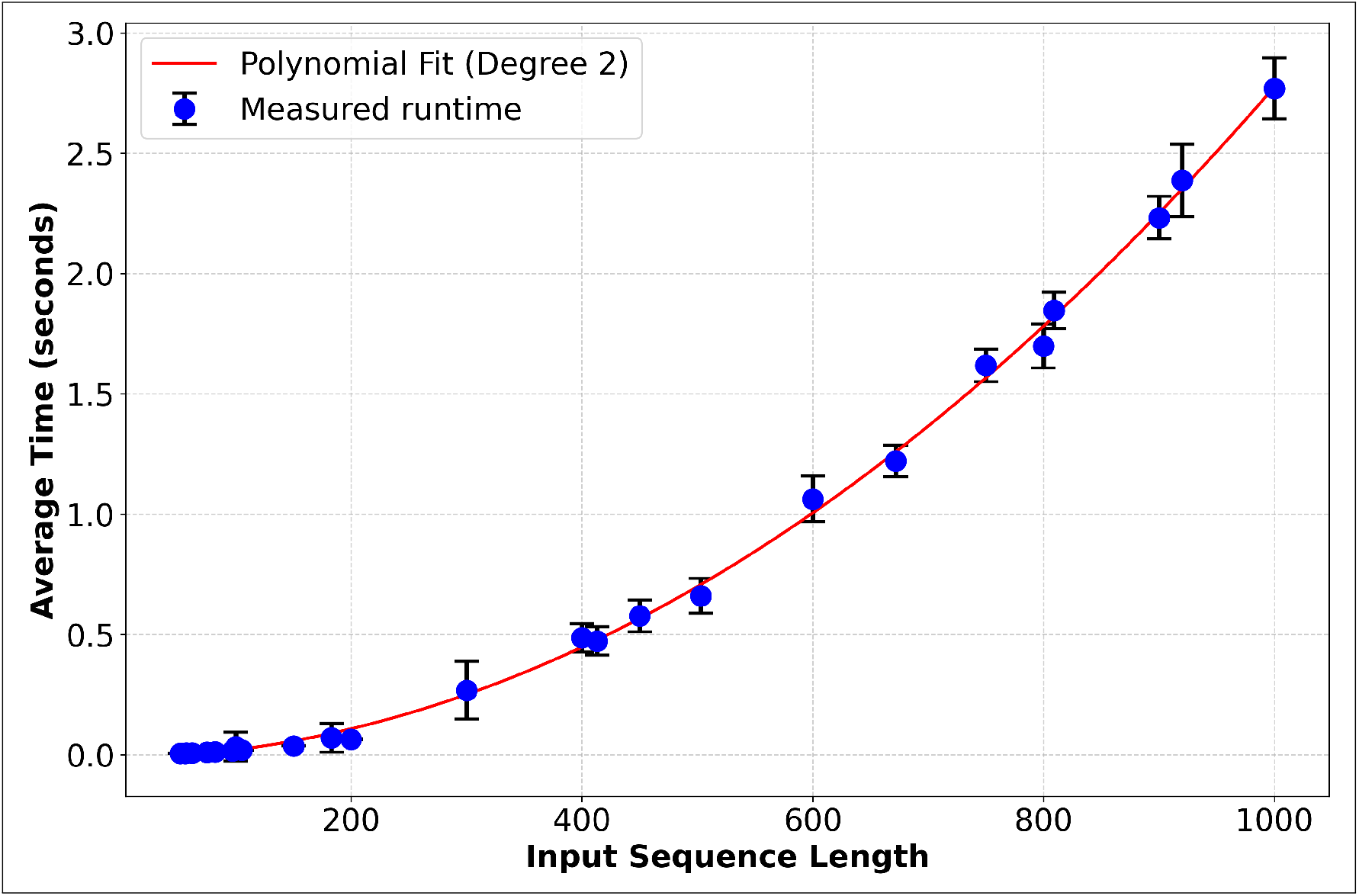
Measurements of the algorithm runtime follows near quadratic scaling with an empirical polynomial degree of 2.068 on Intel(R) Xeon(R) Gold 5118 CPU @ 2.30GHz hardware.

## 4 Discussion

Pool PaRTI is a novel unsupervised approach for assigning importance weights to protein residues and generating general-purpose protein sequence embeddings. It outperforms traditional parameter-free pooling methods across a remarkable range of bioinformatics tasks. By leveraging the internal attention matrices of transformer-based protein language models, Pool PaRTI correctly captures complex relationships between and hierarchy among residues directly from sequence information, without relying on explicit structural or functional data.

Our evaluation across four distinct protein-related tasks—fold class prediction, enzyme classification, sub-cellular localization prediction, and protein-protein interaction prediction—demonstrates the robustness and versatility of Pool PaRTI. These tasks represent diverse and important aspects of protein research. Each task relies on different biological properties and information encoded in protein sequences. SCOP level 2 classification task involves a direct prediction of the protein’s global fold, thereby showcasing the pooling methods’ ability to retain structure-related information. Enzymatic activity prediction relies on the capability to encode and recognize functional motifs, such as active sites and catalytic residues, embedded within sequences. This demonstrates the model’s understanding of sequence-to-function relationships. Protein-protein interaction prediction involves identifying patterns across multiple protein representations, while spatial analysis involves clusters of proteins. These four tests comprehensively evaluate our method’s effectiveness across various protein-related prediction tasks and its ability to make accurate predictions at multiple scales. Pool PaRTI successfully identifies key residues in proteins and generates embeddings that consistently achieve statistically significant improvements in key performance metrics in downstream prediction tasks, Our methodologically consistent approach to retrieval, classifier model development, optimization, and testing confirms that Pool PaRTI’s improvements are both reproducible and robust. This demonstrates that its effectiveness does not rely on serendipitous hyperparameter tuning but is consistently achievable across diverse settings. Further supporting its reliability, our analysis across two different PLMs highlights Pool PaRTI’s effectiveness while also revealing a slight dependence on the underlying model. When paired with ESM2, Pool PaRTI consistently outperforms alternatives across all four tasks. However, with protBERT, its advantage narrows—Pool PaRTI leads in two tasks while remaining competitive in the other two.

It is important to note that prior research has established ESM models, including ESM2 650M and ESM1b, as significantly stronger than protBERT for various downstream tasks such as secondary structure prediction [6], enzyme annotation [22], GO code annotation [23], and binding predictions [24]. This performance gap may stem from protBERT’s smaller size (420 million parameters) compared to ESM2’s 650 million parameters. Pool PaRTI’s effectiveness depends on both high-quality embeddings and high-resolution attention matrices. If protBERT produces lower-resolution attention matrices than ESM2, Pool PaRTI may be at a disadvantage relative to methods that rely only on high-quality token embeddings. Furthermore, because Pool PaRTI normalizes importance scores to sum to 1 before computing weighted average token embeddings, its embedding magnitudes do not inherently correlate with protein size. As a result, Pool PaRTI is better suited for capturing intrinsic, size-independent properties and may be less effective for predicting properties that depend on sequence length. For example, protein connectivity has been shown to correlate with sequence length [25], as longer sequences can accommodate more binding sites. In such a setting, a length-dependent embedding generation method like sum pooling can have an advantage, as demonstrated by its relatively high performance in the protein-protein interaction task.

The SCOP family alignment results suggest that Pool PaRTI preserves more structural and functional information in sequence embeddings compared to baseline methods. Notably, ESMFold has demonstrated its ability to predict protein structures by performing a regression over all attention matrices [2]. Since Pool PaRTI similarly extracts information from attention matrices, its performance gains over other parameter-free sequence embedding methods are consistent with earlier findings in the literature. The preservation of the structural and functional information through the attention matrices was observed not only in ESM2 but also in protBERT, indicating that Pool PaRTI’s performance gain may extend to other self-attention-based protein language model encoders.

One of the most striking features of Pool PaRTI is its interpretability. Our analysis of well-characterized proteins, such as Cytochrome C and superoxide dismutase-1 (SOD1), reveals a strong correlation between residues assigned higher importance weights by Pool PaRTI and known functionally or structurally significant regions. This emergent property, achieved without explicit training on structural or functional data, suggests that Pool PaRTI captures intrinsic sequence features that correlate with functional importance, offering new insights into protein structure-function relationships.

Traditional pooling methods, such as mean pooling, max pooling, and sum pooling lack this interpretability. These approaches aggregate token embeddings into a single vector without providing any indication of which sequence regions contribute most significantly to the final representation. In contrast, Pool PaRTI’s assignment of importance weights to individual residues allows for a more nuanced understanding of how different regions of a protein sequence influence the overall representation. This interpretability feature of Pool PaRTI opens up new avenues for protein analysis. It may aid in the identification of potential functional sites, guide experimental design for mutagenesis studies, or provide valuable insights into protein engineering applications. Furthermore, the ability to highlight potentially important regions without relying on structural data makes Pool PaRTI particularly valuable for studying proteins with unknown structures or intrinsically disordered regions.

The general-purpose nature of Pool PaRTI embeddings is a key advantage. Unlike methods that require task-specific training or optimization, our approach generates informative sequence embeddings without the need for task-specific adjustments. By employing PageRank instead of more complex graph learning algorithms, we deliberately avoid parameter optimization tailored to specific downstream tasks. This choice enables Pool PaRTI to create general-purpose token importance weights, enhancing its versatility across various protein analysis tasks. This design makes Pool PaRTI particularly valuable for exploratory analyses and for applications in data-scarce domains where task-specific training data may be limited for additional pooling parameter optimization. Pool PaRTI operates independently from the neural network’s training phases, allowing it to be integrated with any self-attention-based protein language model. This feature facilitates its application without the need for additional training or alterations to the original model architecture, further reinforcing its task-agnostic nature.

From a computational perspective, our analysis shows that Pool PaRTI does not introduce prohibitive overhead compared to other pooling methods, particularly for sequence lengths relevant to the human proteome. This efficiency, coupled with its superior performance in supporting downstream tasks, makes Pool PaRTI a practical choice for large-scale protein analysis research.

There are several limitations to Pool PaRTI. First, we primarily tested the method using two encoder-only, representation-focused protein language models. While the use of *NxN* sized self-attention matrices theoretically enables application to other transformer-based models, we have not explicitly demonstrated its effectiveness beyond these PLMs. Adapting Pool PaRTI to generative decoder models—where attention is autoregressive and directional—and to attention-free state-space models falls outside the scope of this work. Second, although our analysis shows that Pool PaRTI is computationally efficient for typical protein sequence lengths, its performance for extremely long outlier sequences may warrant further investigation. While Pool PaRTI offers interpretability unlike the traditional pooling methods, the biological significance of the assigned importance weights may require careful analysis in some contexts, especially for proteins with multiple modes of interactions. Lastly, we have not demonstrated the generalizability of Pool PaRTI in other non-protein sequence analysis domains, such as natural language processing. This remains a theoretical possibility that has not yet been empirically tested.

## 5 Conclusion

We introduced Pool PaRTI, a novel approach to generating protein sequence embedding from individual token embeddings. By combining attention matrices from transformer-based protein language models with the PageRank algorithm, Pool PaRTI captures the complex hierarchy of residues within protein sequences. It assigns importance weights to residues based on their global context, creating sequence embeddings that reflect both local and global interactions. This approach effectively distills the rich information from token-level embeddings into an expressive sequence representation.

## 7 Data and code availability

Our PyTorch implementation of Pool PaRTI and the experimental task models are available at https://github.com/Helix-Research-Lab/Pool_PaRTI.git. The Pool PaRTI residue importance values for all human proteins on UniProt are available at https://zenodo.org/records/15036725 for ESM2 and protBERT. The data we used in these tasks are as follows: For the subcellular localization prediction task, we used the data annotation and splits provided by Thumuluri et al [7]. For the protein-protein interaction prediction tasks, we used the data annotation and gold standard split provided by Bernett et al. [21]. For the enzyme classification task, we used the gold standard ECPred40 dataset created by Buton et al. [19]. We obtained the SCOP labels from the SCOP database [17, 18] with API access.

## 8 Acknowledgements

We thank the Altman Lab members and Dr. Brian Hie for useful discussions. We also thank Chetan Nair and Hee Jung Choi for helping explore some use cases outside biology. We performed the method development and analysis on the Sherlock cluster, and we thank Stanford University and Stanford Research Computing Center for the computational resources. A.T is supported by Smith Fellowship (Stanford Graduate Fellowship), G.N. is supported by NLM Training Grant 5T15007033-40 and NIH NLM 1F31LM014646. R.B.A. is supported by NIH GM102365, NIH GM153195, and Chan Zuckerberg Biohub.

## 10 Funding

Smith Fellowship (Stanford Graduate Fellowship)

NLM Training Grant 5T15007033-40

NIH NLM 1F31LM014646

NIH GM102365

NIH GM153195

Chan Zuckerberg Biohub

## A Appendix / Supplemental Material

### A.1 Theoretical expectation and standard deviation of Jaccard index under random weight assignment

In this section, we derive the expectation and standard deviation of the Jaccard index when two sets of indices are selected randomly from a total of *N* elements.

#### A.1.1 Expectation of the Jaccard Index

Let *p* = *k/N* be the fraction of selected elements, where *k* is the number of selected elements from a total of *N* . The Jaccard index for two randomly selected sets of size *k* is given by:

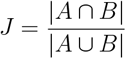

where *A* and *B* are the two selected sets. The key quantities to compute are:

- **Expected size of the intersection:** Since each element is included in a set with probability *p*, the probability that a given element appears in both sets is *p*^2^. Since there are *N* elements, the expected intersection size is:

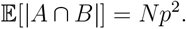
- **Expected size of the union:** Using the inclusion-exclusion principle, the expected union size is:

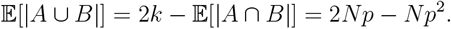

Thus, the expectation of the Jaccard index is:

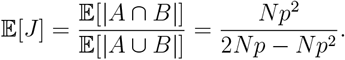

Simplifying,

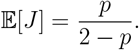

This is the expected Jaccard index for randomly sampled sets as a function of *p*, independent of *N* .

#### A.1.2 Variance and Standard Deviation of the Jaccard Index

To compute the variance of *J*, we use the formula:

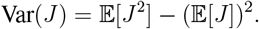

Following standard combinatorial derivations from random set theory, the variance of *J* is:

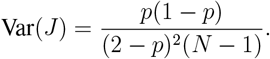

Taking the square root, the standard deviation of *J* is:

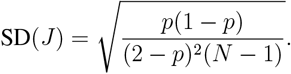

### A.2 The choice of metrics

For evaluation of the performance gains that the Pool PaRTI embeddings yield, we use the Matthews Correlation Coefficient (MCC) and Area Under the Precision-Recall Curve (AUPRC), chosen for their robustness in assessing binary classification performance across unbalanced datasets, alongside accuracy. For multi-label prediction, we also report the Jaccard index to evaluate performances. For single label and multilabel binary classifications, we employed different metrics to assess model predictive performances under the influence of the input embeddings resulting from alternative pooling methods.

#### A.2.1 Precision @10%

Precision at 10% (P@10%) is a useful metric for ranking-based tasks, measuring how many of the top 10% of predictions are correct. In this context, 10% refers to the 10% of the size of each category in balanced subsampling. For example, if we sample a balanced subset from an imbalanced dataset such that each category has 30 representatives, we report the precision for the labels of the top 3 proteins that have the highest cosine similarity to the query protein, based on the label of the query protein. This is particularly relevant for applications where the model needs to rank the most relevant items higher. It is defined as:

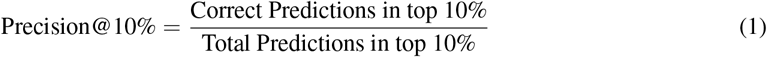

This metric helps evaluate the effectiveness of embeddings in ranking scenarios and ensures that highly relevant items are correctly prioritized [26].

#### A.2.2 Mean Reciprocal Rank (MRR)

The Mean Reciprocal Rank (MRR) is a ranking evaluation metric that computes the average of the reciprocal ranks of the first correct prediction. It is particularly useful in search systems, where finding the first relevant result is critical. MRR is computed as follows:

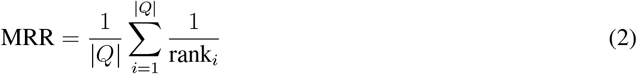

where is the number of queries and is the rank position of the first relevant item for the -th query.

Higher MRR values indicate better performance in ranking tasks, as relevant predictions appear earlier in the ranked list [26].

#### A.2.3 Jaccard Index

The Jaccard Index, also known as Jaccard similarity, is a measure used to compare the similarity between two sets. It is particularly useful in multi-label classification problems, where it measures the overlap between predicted and true label sets. It is defined as:

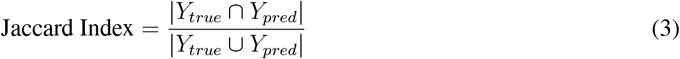

where is the set of true labels and is the set of predicted labels.

The Jaccard Index ranges from 0 to 1, with higher values indicating greater similarity between the predicted and actual labels. It is particularly effective in evaluating model performance in multi-label scenarios, ensuring that both precision and recall are considered simultaneously [26].

#### A.2.4 Accuracy

Accuracy is the simplest and most commonly used metric for evaluating classification models. It is defined as the ratio of correctly predicted instances to the total instances. Mathematically, accuracy is expressed as:

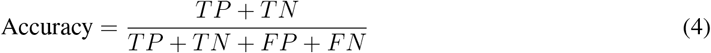

While accuracy is intuitive and easy to understand, it can be misleading for imbalanced datasets, as it does not differentiate between the types of errors (false positives and false negatives).

#### A.2.5 Balanced Accuracy for Multicategory Predictions

Balanced accuracy is an extension of accuracy that accounts for class imbalance in multi-category classification problems. Unlike standard accuracy, which can be skewed by dominant classes, balanced accuracy ensures that each class contributes equally to the overall performance. It is computed as the average recall across all classes:

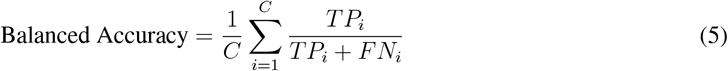

where *C* is the total number of classes, and *TP*_*i*_ and *FN*_*i*_ represent the true positives and false negatives for class *i*, respectively.

Balanced accuracy is particularly useful in scenarios where some categories are underrepresented, preventing models from favoring majority classes. In a balanced dataset, balanced accuracy is equal to the standard definition of accuracy. It provides a more fair assessment of performance across all classes and is widely used in multi-class classification tasks [27].

#### A.2.6 Matthews Correlation Coefficient (MCC)

The Matthews Correlation Coefficient (MCC) is a measure of the quality of binary (two-class) classifications. It takes into account true and false positives and negatives and is generally regarded as a balanced measure which can be used even if the classes are of very different sizes. The MCC is defined as:

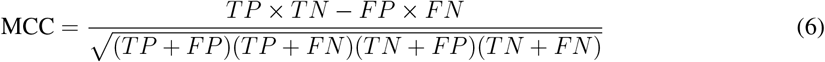

where:

- *TP* = True Positives
- *TN* = True Negatives
- *FP* = False Positives
- *FN* = False Negatives

MCC returns a value between -1 and 1. A coefficient of 1 represents a perfect prediction, 0 represents a random prediction, and -1 indicates total disagreement between prediction and observation [28].

#### A.2.7 Area Under the Precision-Recall Curve (AUPRC)

The Area Under the Precision-Recall Curve (AUPRC) is a performance measurement for classification problems at various threshold settings, especially useful for imbalanced datasets. The Precision-Recall curve plots precision (positive predictive value) against recall (sensitivity) for different threshold values. Precision and recall are defined as:

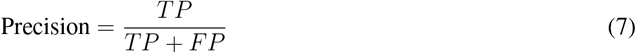

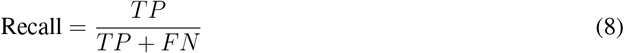

The AUPRC provides a single scalar value to summarize the curve, with higher values indicating better performance. It is particularly useful when the positive class is rare, as it focuses on the performance of the positive class [29].

These metrics together provide a comprehensive evaluation framework for our models, allowing us to capture different aspects of performance, particularly in the presence of class imbalance and multi-label scenarios.

### A.3 Empirical analysis of Pool PaRTI runtime scaling

To formally assess the empirical scaling of runtime for the Pool PaRTI algorithm, we executed the algorithm on 25 protein sequences, repeating each run 15 times. We then applied a logarithmic transformation to both the runtime data and the corresponding sequence lengths. A linear regression model was fitted to the log-transformed data, yielding a slope of 2.068, which aligns with our theoretical reasoning in Section 2. This slope indicates the exponent in the power-law relationship between the algorithm’s runtime and the input size, providing a quantitative measure of the algorithm’s scaling behavior with respect to input size.

### A.4 Deep Learning Models and Configurations

#### A.4.1 Subcellular Localization Prediction Task

Description of the model

- 2 x (linear layer + leaky ReLU + dropout)
- Linear layer
- Input residual connection
- Sigmoid Fixed configurations
- Initial learning rate: 0.01
- Max number of epochs: 1000
- Early stopping patience epochs: 50
- Optimizer: AdamW
- Learning rate scheduler: ReduceLROnPlateau
- Optimizer learning rate patience epochs: 10
- Learning rate reduction ratio: 0.1
- Weight Initialization: Xavier normal
- Random seed: 42
- Gradient clip value: 5.0
- batch size: 64 Hyperparameter optimization space
- Weight decay: [0.01, 0.1, 0.2]
- Dropout rate: [0.15, 0.25]
- Slope of Leaky ReLU: [0.01, 0.1]
- Exponent for imbalance penalty taming: [0.75, 1, 1.25]

#### A.4.2 Protein-Protein Interaction Prediction Task

Description of the model

- 1 x (linear layer + batch norm + leaky ReLU + dropout)
- 1 x (linear layer + linearly transformed residual connection + leaky ReLU)
- Linear layer Fixed configurations
- Hidden dimensions: 1024
- Max number of epochs: 40
- Early stopping patience: 9
- Optimizer: AdamW
- Learning rate scheduler: ReduceLROnPlateau
- Optimizer learning rate patience epochs: 4
- Random seed: 42
- Weight initialization: Kaiming normal
- Gradient clip max value: 2.0
- batch size: 32 Hyperparameter optimization space
- Initial learning rate: [0.001, 0.01]
- Weight decay: [0.01, 0.1, 0.2]
- Dropout: [0.05, 0.15, 0.25]
- Reduction in learning rate ratio: [0.2, 0.5]
- Leaky ReLU slope: [0.01, 0.1, 0.2]

### A.5 Compute resources for experiments

We ran all computational experiments and pooling algorithms on Tesla V100-SXM2-16GB GPUs housed in the internal Sherlock cluster. For each task, the hyperparameter optimization experiments were limited to two days on GPU. On the same GPUs, we conducted unpublished preliminary experiments in developing the Pool PaRTI algorithm. We generated the precomputed ESM2 token embeddings on NVIDIA A100-PCIE-40GB GPUs, also housed in the internal Sherlock cluster. Token embedding generation took 70 GPU hours through the ESM2 650M model and 50 hours through protBERT model. We have computed token embeddings once for each sequence and performed several different pooling operations on the precomputed token embeddings before feeding the sequence embeddings as inputs to the respective models.

